# Tendon and Motor Phenotypes in the *Crtap^-/-^* Mouse Model of Recessive Osteogenesis Imperfecta

**DOI:** 10.1101/2020.04.21.048488

**Authors:** Matthew W. Grol, Nele A. Haelterman, Joohyun Lim, Elda M. Munivez, Marilyn Archer, David M. Hudson, Sara F. Tufa, Douglas R. Keene, Kevin Lei, Dongsu Park, David R. Eyre, Brendan H. Lee

## Abstract

Osteogenesis imperfecta (OI) is characterized by short stature, skeletal deformities, low bone mass with bone fragility, and motor deficits. A subset of OI patients also present with joint hypermobility; however, the role of tendon/ligament dysfunction in OI pathogenesis is largely unknown. Using the *Crtap*^-/-^ mouse model of severe, recessive OI, we found that mutant Achilles tendons and patellar ligaments were thinner with increased collagen cross-links and reduced collagen fibril size at 1- and 4-months compared to wildtype. Patellar ligaments from *Crtap*^-/-^ mice also had fewer progenitors with a concomitant increase in immature cells. RNA-seq analysis of Achilles tendons and patellar ligaments from 1-month *Crtap*^-/-^ mice revealed dysregulation in matrix gene expression concomitant with predicted alterations in TGF-β, inflammatory, and metabolic signaling. Finally, a series of behavioral tests revealed severe motor impairments and reduced grip strength in 4-month *Crtap*^-/-^ mice – a phenotype that correlates with the tendon/ligament pathology.

## INTRODUCTION

Tendon is a fibrous tissue that connects skeletal muscle to bone to facilitate motion, whereas ligaments connect articulating bones to support joint alignment and function^1, 2^. The extracellular matrix of tendons/ligaments is primarily composed of type I collagen as well as smaller quantities of other collagens and proteoglycans^3^. During development, the collagen fibrils in tendons and ligaments develop through addition and lengthening before transitioning to the appositional fusion of existing fibers with continued lengthening in postnatal life^4^. The synthesis and assembly of this collagen-rich matrix are influenced by other small collagens and proteoglycans as well as by the cross-linking chemistry of type I procollagen fibrils, which in turn regulates fibril size and strength^5^. Like tendon and ligament, the organic matrix of bone consists largely of type I collagen^6^, and disruptions in collagen synthesis and folding are known to negatively impact its biochemical and mechanical properties in connective tissue diseases such as Osteogenesis Imperfecta (OI)^7^. However, despite evidence of joint mobility phenotypes and motor deficits in OI patients^8, 9^, tendon and ligament phenotypes in this disease are relatively understudied.

OI is a heterogeneous group of disorders characterized by variable short stature, skeletal deformities, low bone mass, and increased bone fragility. Approximately 80% of OI cases are caused by dominantly inherited mutations in the genes encoding the α1(I) or α2(I) chains of type I collagen. Mutations in genes responsible for the synthesis, post-translational modification, and processing of collagen, such as cartilage-associated protein (CRTAP), lead to severe, recessive forms of this disease^7^. In addition to skeletal defects, other connective tissue manifestations including joint hypermobility and skin hyperlaxity are observed in a subset of OI patients^8, 9^. Our and others’ studies have shown that CRTAP forms a complex with Prolyl 3-hydroxylase 1 (P3h1, encoded by *Lepre1*) and Cyclophilin B (CypB, encoded by *Ppib*), and is required for prolyl 3-hydroxylation of type I procollagen at Pro986 of chain α1(I) and Pro707 of chain α2(I)^10–12^. In this regard, loss of either CRTAP or P3H1 leads to loss of this complex and its activity, causing a severe recessive form of OI characterized by short stature and brittle bones^11, 13–15^. Collagen isolated from *Crtap^-/-^* and *Lepre1^-/-^* mice is characterized by lysine over-modifications and abnormal fibril diameter^11, 14^. While a comprehensive analysis of *Crtap*^-/-^ mice has revealed multiple connective tissue abnormalities, including in lungs, kidneys, and skin^16^, the impact of the loss of Crtap on tendons and ligaments remains unknown.

Alterations in collagen fibril size and cross-linking have been noted in a limited number of studies using dominant or recessive mouse models of OI^17–19^; however, whether loss of Crtap impacts tendon and ligament development and structure remains unknown. In this study, we show that *Crtap*^-/-^ mice have thinner Achilles tendons and patellar ligaments at 1 and 4 months-of-age that are hypercellular with a reduction in tendon progenitors and total ligament volume. Moreover, the patellar and cruciate ligaments from *Crtap*^-/-^ show increased cell size and variable ectopic chondrogenesis by 4 months-of-age. Examining the collagen matrix, we found an increase in stable (irreversible) collagen cross-links at both timepoints, accompanied by alterations in fibril diameter at 4-months compared to wildtype controls. RNA-seq analyses revealed both common and distinct changes in the transcriptome of the Achilles tendon and patellar ligaments of *Crtap*^-/-^ mice compared to wildtype with a predicted activation of transforming growth factor-β (TGF-β) and inflammatory signaling in both tissues. These changes in *Crtap*^-/-^ tendons and ligaments were accompanied by motor deficits and reduced strength at 4 months-of-age. In conclusion, loss of CRTAP in mice causes tendon/ligament abnormalities and significant behavioral impairments.

## MATERIALS AND METHODS

### Animals

*Crtap*^-/-^ mice were generated as previously described^11^ and maintained on a mixed C57BL/6J and 129Sv genetic background. All studies were performed with approval from the Institutional Animal Care and Use Committee (IACUC) at Baylor College of Medicine. Mice were housed 3 to 4 mice to a cage in a pathogen-free environment with *ad libitum* access to food and water and under a 14h light/10h dark cycle.

### Histological analysis

Mice were euthanized and ankle and knee joints were dissected and fixed for 48 h on a shaker at 4°C or room temperature in freshly prepared 4% paraformaldehyde (PFA) in 1×phosphate-buffered saline (PBS). Samples were decalcified at 4°C using 10% ethylenediaminetetraacetic acid (EDTA) in 1×PBS for 10 days (with one change out at 5 days) before paraffin embedding using a standard protocol. Samples were sectioned at 6-μm and stained with hematoxylin and eosin (H&E) to visualize tendon structures. Cell number per tissue area was determined using the Fiji release of ImageJ^20^.

### Fluorescence-activated cell sorting analysis of tendon progenitors

After dissection of patellar ligaments from 5-month-old wildtype and *Crtap*^-/-^ mice, tissues were cut into small pieces in 1×PBS with 10% fetal bovine serum (FBS) and incubated with 500 μl of PBS + 10% FBS and 0.1% collagenase at 37°C for 3 hours. After digestion, cells were filtered with a 40-μm strainer, washed, resuspended in 1×PBS at a concentration of 10^6^ cells/mL, and stained with CD45-pacific blue (clone: 30-F11), CD31-eFlour 450 (clone: 390), CD146-PE-Cy7 (clone ME-9F1) and CD200-APC (clone OX-90) (eBioscience). Propidium iodide was used for selecting viable cells. Cell analysis was performed using a LSRII Fortessa, and FACS experiments were done using an AriaII cytometer (BD Biosciences, San Jose, CA). Data were analyzed with FlowJo software (TreeStar, OR).

### Phase-contrast μCT imaging and analysis

To quantify tendon/ligament volume, knee joints were dissected from mice, stained with contrast agents, scanned by phase-contrast μCT, and analyzed using TriBON software (RATOC, Tokyo, Japan) as previously described for articular cartilage^21–24^. In addition to the articular surfaces (data not shown), we performed contrast-enhanced visualization of the patellar ligament using this protocol. To quantify volume, samples were examined in transverse where the patellar ligament boundary was easily distinguished from the joint capsule. Ligament volume was assessed from its origin within the patella to its insertion at the tibia.

### Transmission electron microscopy analysis of collagen fibril size

Mouse ankle and knee joints were dissected and fixed in fresh 1.5% glutaraldehyde/1.5% PFA (Tousimis) with 0.05% tannic acid (Sigma) in 1×PBS at 4°C overnight to preserve the native tension on relevant tendons/ligaments. The next day, flexor digitorum longus (FDL) and Achilles tendons as well as patellar ligament were dissected out in 1×PBS, and placed back into fixative. Samples were then post-fixed in 1% osmium tetroxide (OsO_4_), rinsed in Dulbecco’s Modified Eagle Medium (DMEM), and dehydrated in a graded series of ethanol to 100%. Samples were rinsed in propylene oxide, infiltrated in Spurrs epoxy, and polymerized at 70°C overnight. TEM images were acquired using an FEI G20 TEM at multiple magnifications to visualize transverse sections of collagen fibrils. Collagen fibril diameter was measured using the Fiji release of ImageJ^20^.

### Tendon collagen cross-linking analysis

Collagen hydroxylysyl-pyridinoline (HP) cross-links were quantified as previously described^25, 26^. In brief, tendons and ligaments isolated from hindlimbs were hydrolyzed by 6M HCl for 24h at 108°C. Dried samples were then dissolved in 1% (v/v) n-heptafluorobutyric acid for quantitation of HP by reverse-phase HPLC with fluorescence monitoring.

### RNA-seq analysis

At 1 month-of-age, Achilles tendons were excised using scissors proximal to the calcaneal insertion and distal to the tendon-muscular-junction, whereas patellar ligaments were removed via scalpel just proximal to the tibial insertion and distal to the patella. Tendons and ligaments were not cleaned of their paratenon layers in either case. RNA extraction for both tissues was performed with the Fibrous Connective Tissue kit (Qiagen), with columns from the RNeasy Micro Kit (Qiagen) employed for the patellar ligament. Quality control, library preparation, sequencing, and differential gene expression analysis including gene ontology analysis was performed by GENEWIZ^®^ (South Plainfield, NJ). For the examination of predicted Upstream Regulators, we utilized the Ingenuity Pathway Analysis (IPA) platform (Qiagen) with an adjusted *p*-value of 0.05. The expression is shown as Base Means and read counts of genes were normalized per million transcripts (Transcripts Per Million; TPM).

### Open field assessment of spontaneous motor activity

Open field activity was measured using the VersaMax Animal Activity Monitoring System (AccuScan Instruments, Columbus, OH). On the day of assessment, mice were transferred to the test room and allowed to acclimate in their home cage for 30 min at 50 Lux of illumination with 60 dB of white noise. Mice were then placed individually into clear 40 cm × 40 cm × 30 cm chambers and allowed to move freely for 30 min. Locomotion parameters and zones were recorded using the VersaMax activity monitoring software. Chambers were cleaned with 30-50% ethanol to remove the scent of previously tested mice between each run.

### Rotarod analysis of motor coordination and endurance

On the day of assessment, mice were transferred to the test room and allowed to acclimate within their home cage for 30 min at 50 Lux of illumination with 60 dB of white noise. Mice were then placed on a rotarod (UGO Basile, Varese, Italy) set to accelerate from 5-to-40 RPM over 5 min. Five trials were performed per day for 2 consecutive days (trials 1-10) with a rest time of 5 min between trials. Latency to fall was recorded when the mouse fell from the rotating rod or went for two revolutions without regaining control. The rotarod was cleaned with 30-50% ethanol between mice to remove the scent of previously tested animals.

### Grid foot slip analysis of motor coordination

The grid foot slip assay consisted of a wire grid set atop a stand where the movement was recorded by a suspended digital camera. Mice were transferred to the test room on the day of assessment and allowed to acclimate within their home cage for 30 min at 50 Lux of illumination with 60 dB of white noise. Mice were then placed one at a time on the grid and allowed to move freely for 5 min. The observer sat 6-8 feet away at eye-level to the mouse and recorded forelimb and hindlimb foot slips using the ANY-maze video tracking software (Stoelting Co., Wood Dale, IL). After the test, mice were removed to their original home cage. Forelimb and hindlimb foot slips were normalized to the total distance traveled during the test.

### Inverted grid analysis of strength and endurance

On the day of assessment, mice were transferred to the test room and allowed to acclimate within their home cage for 30 min at 50 Lux of illumination with 60 dB of white noise. Mice were then placed in the middle of a wire grid, held approximately 18-in above a cushioned pad, and inverted. The latency to fall for each mouse was recorded. At the completion, mice were returned to their home cage.

### Grip strength analysis

Mice were transferred to the test room on the day of assessment and allowed to acclimate within their home cage for 30 min at 50 Lux of illumination with 60 dB of white noise. Each mouse was then lifted by its tail onto the bar of a digital grip strength meter (Columbus Instruments, Columbus, OH). Once both forepaws had gripped the bar, the mouse was gently pulled away from the meter by its tail at a constant speed until the forepaws were released. The grip (in N of force) was recorded and the procedure repeated twice for a total of three measurements, which were averaged for the final result.

### Statistical analysis

Determination of sample size was based on previous publications. Biological replicates were defined as an individual mouse for each experiment. Respective tendons and ligaments from the left and right hindlimbs were combined for TEM, collagen cross-linking, tendon progenitor, and RNA-seq analyses. Data are presented as means ± S.D. or min-to-max box and whisker plots with individual data points. For data whose residuals passed the Shapiro-Wilk test for normality, groups of two were compared using unpaired t-tests, and groups of three or more were compared using one-way ANOVA followed by Tukey’s post-hoc tests. For data whose residuals did not have a normal distribution, groups of two were compared using a Mann-Whitney test, and groups of three or more were compared using Kruskal-Wallis followed by Dunn’s post-hoc tests. For the Rotarod assay, where time was a variable, groups were compared using repeated measures two-way ANOVA followed by Tukey’s post-hoc tests. For all tests reported above, statistical analysis was performed using Prism 8.3.1 (GraphPad Software, La Jolla, CA). For all tests, the exact *p*-value is reported, and a *p*-value of < 0.05 was considered statistically significant.

## RESULTS

### Crtap^-/-^ mice exhibit abnormal tendon development

Mice lacking Crtap present with growth delay, rhizomelia, and severe osteoporosis together with disruption of other connective tissues including lung and skin^11, 16^. To assess whether *Crtap^-/-^* mice exhibit disruptions in the development of tendons and ligaments, we harvested ankle and knee joints at 1 and 4 months-of-age to histologically examine the Achilles tendon and patellar ligament. At 1-month, *Crtap^-/-^* mice presented with thinner Achilles tendons and patellar ligaments (**Figure 1C,F**) compared to wildtype (**Figure 1A,D**) and heterozygous mice (**Figure 1B,E**) with increased cell density in both structures (**Figure 1M**). There was also prominent hyperplasia of the synovial membrane in *Crtap* mutant knees at this age (**Figure 1F**). By 4 months-of-age, *Crtap^-/-^* Achilles tendons and patellar ligaments remained thinner and hypercellular compared to wildtype and heterozygous mice (**Figure 1G-L,N**). In addition, synovial hyperplasia persisted within *Crtap*^-/-^ knees (**Figure 1L**). Interestingly, ectopic chondrogenesis was present towards either end of the patellar ligament and/or cruciate ligaments in some (but not all) 4-month-old *Crtap^-/-^* mice (**Figure 1L**) – a phenomenon that can occur in tendinopathy^27^.

**Figure 1.**
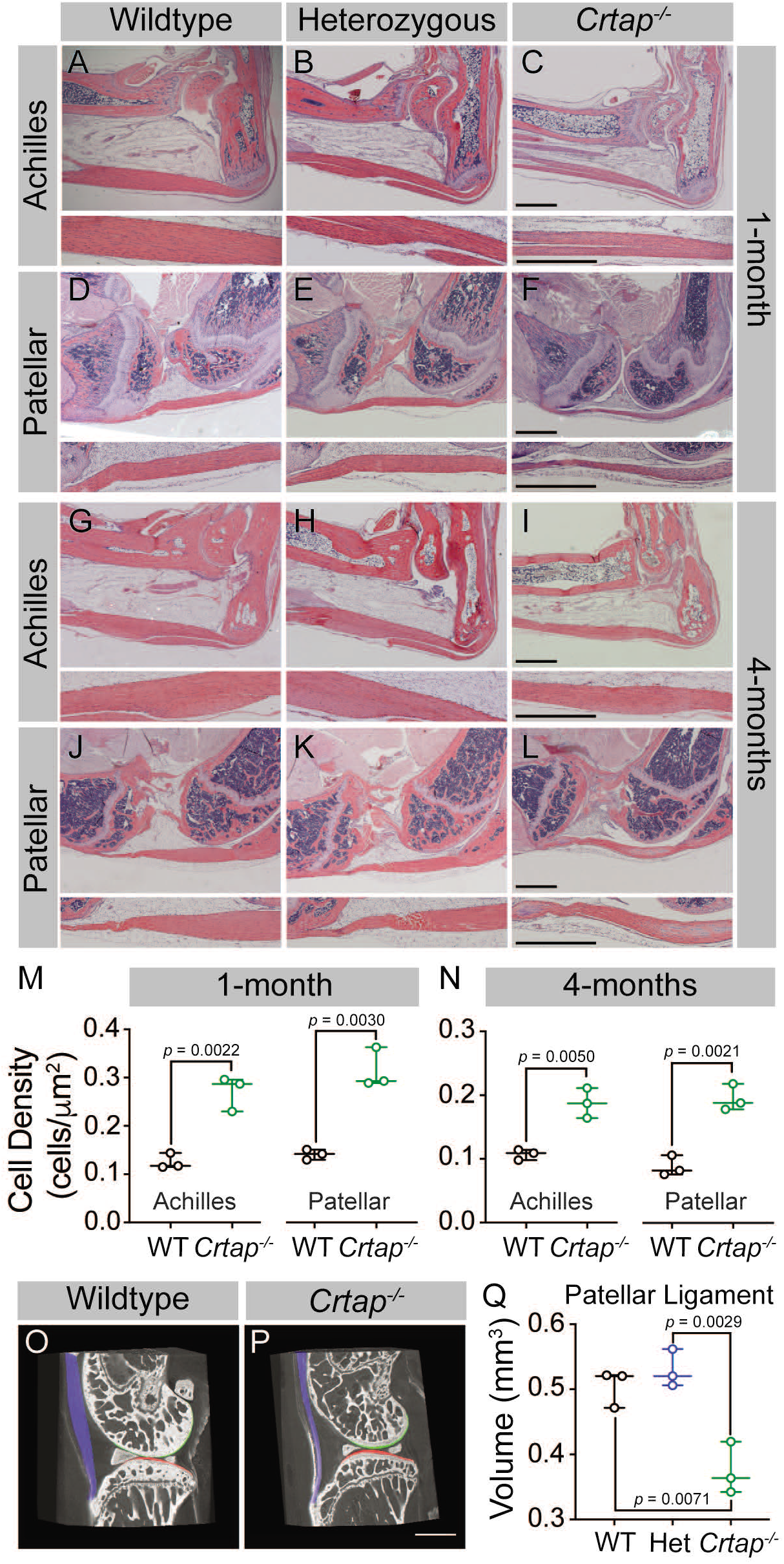
Loss of Crtap causes thinning and hypercellularity of tendons and ligaments in young and mature mice. **(A-C)** representative H&E images of 1-month ankle joints. **(D-F)**, representative H&E images of 1-month knee joints. **(G-I)** representative H&E images of 4-month ankle joints. **(J-L)** representative H&E images of 4-month knee joints. For all micrographs, higher magnification images of the mid-tendon/ligament are illustrated. n = 3-4 mice per group. Scale bar is 1-mm. **(M-N)** quantification of cell density for Achilles tendons and patellar ligaments of wildtype and *Crtap^-/-^* mice at 1-month **(M)** and 4-months **(N)** of age. Data are min-to-max box and whisker plots with individual points indicated. n = 3 mice per group. Data passed the Shapiro-Wilk test for normality, and groups were compared using two-tailed unpaired t-tests. Exact *p*-values are reported. **(O-P)** representative phase-contrast μCT images of 4-month wildtype **(O)** and *Crtap^-/-^* **(P)** knee joints. Blue indicates the patellar ligament, green indicates the femoral articular cartilage, and red indicates the tibial articular cartilage. Scale bar is 1-mm. **(Q)** volumetric quantification of the patellar ligament volume in wildtype, heterozygous, and *Crtap^-/-^* mice. Data are min-to-max box and whisker plots with individual points indicated. n = 3 mice per group. Data passed the Shapiro-Wilk test for normality, and groups were compared using one-way ANOVA with Tukey’s post-hoc tests. Exact *p*-values are reported.

To more accurately quantify the differences in tendon/ligament size between genotypes, we performed phase-contrast μCT to examine differences in patellar ligament volume between genotypes (**Figure 1O-Q**) – a technique we previously used to examine soft tissues including articular cartilage in intact murine knee joints^21–24^. In 4-month-old mice, we observed a thinning of the patellar ligament compared to wildtype mice, with no notable changes in articular cartilage surface (**Figure 1O,P**). Confirming our histological findings, quantification revealed a decrease in patellar ligament volume in *Crtap^-/-^*, but not in heterozygous mice, compared to wildtype controls (**Figure 1Q**).

Postnatal progenitor populations are thought to be critical for the maintenance and regeneration of tendon/ligament tissue. Previous studies have shown that a subset of fibroblasts in the tendon midsubstance (endotenon) has the potential to contribute to tendon homeostasis and regeneration at various developmental stages^28–30^. In addition, platelet-derived growth factor receptor a (PDGFRa)^+^ cells in the tendon sheath (paratenon) can contribute to tendon regeneration after acute injury^31^. These progenitor cells can be labeled by *Tppp3*-CreER^31^ or *aSMA*-CreER^28^ with the expression of tendon progenitor markers such as CD146^32^, CD140a (PDGFRa)^31^, and CD200^33^. Given the significant hypercellularity seen in the Achilles tendons and patellar ligaments from *Crtap^-/-^* mice, we examined whether the postnatal progenitor population is altered compared to wildtype controls. Using fluorescence-activated cell sorting (FACS) analysis of 5-month-old patellar ligaments, we observed a significant decrease in the percentage of CD45^-^CD31^-^CD146^+^CD200^+^ stem/progenitor cells (~2%) compared to wild-type mice (~4%) (**Figure 2A-C**, red box), while CD45^-^CD31^-^CD146^-^CD200^+^ immature cells were concomitantly increased in *Crtap^-/-^* mice (**Figure 2D**).

**Figure 2.**
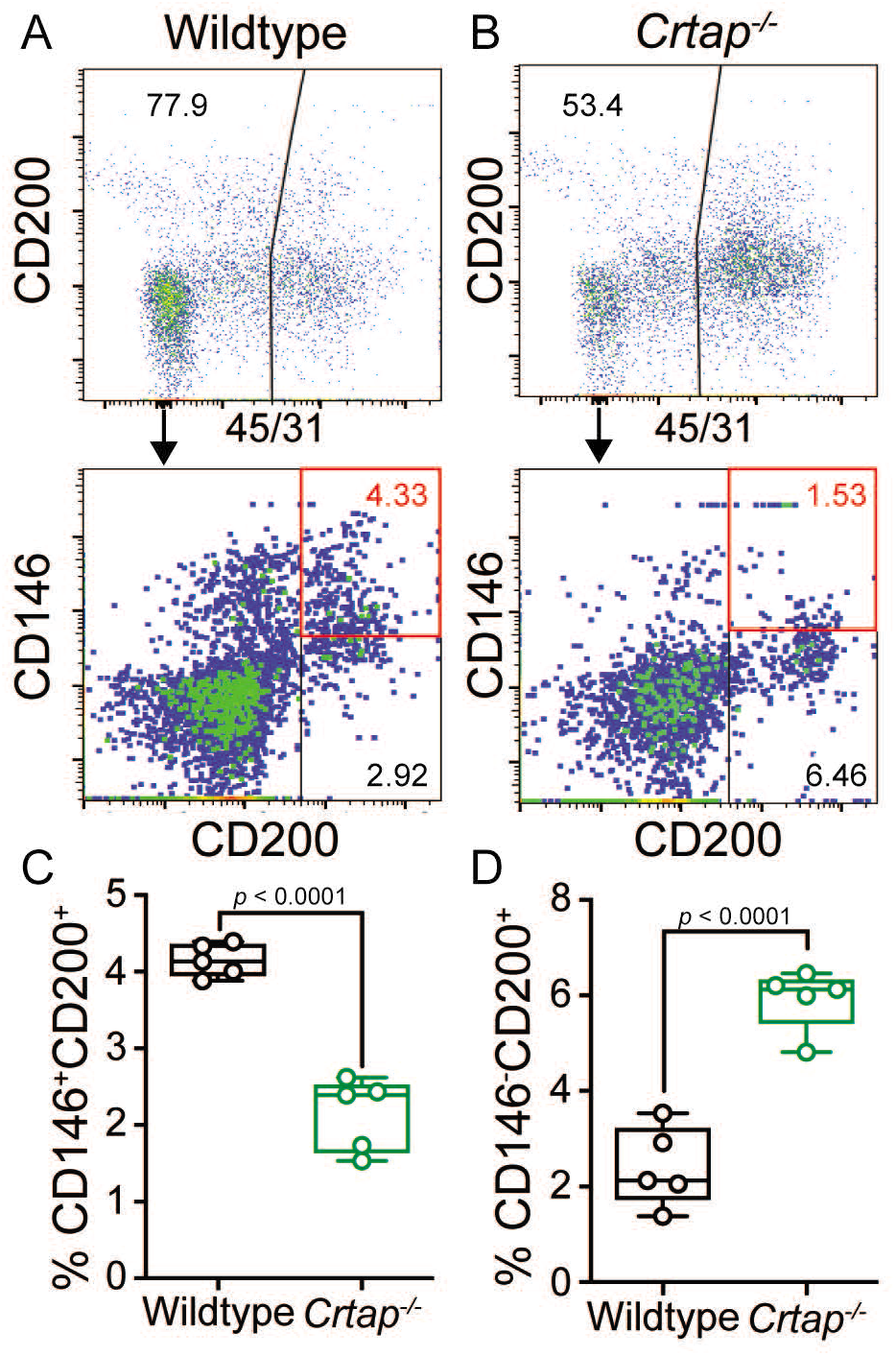
Loss of Crtap in patellar ligaments leads to a decrease in progenitor cells and an accumulation of immature resident tissue cells. **(A-B)** patellar ligament cells isolated from 5-month-old wildtype **(A)** or *Crtap^-/-^* **(B)** mice were analyzed for the expression of CD200 and CD146 tendon/ligament progenitor markers (*top histogram*) within the CD45^-^CD31^-^ population (*bottom histogram*). The plots are representative from a single wildtype or *Crtap^-/-^* mouse. **(C-D)** graphs show the percentage of CD45^-^CD31^-^CD146^+^CD200^+^ progenitor cells **(C)** and CD45^-^ CD31^-^CD146^-^CD200^+^ immature ligament cells **(D)** from 5-month wildtype and *Crtap^-/-^* patellar ligaments. Data are min-to-max box and whisker plots with individual points indicated. n = 5 mice per group. Data passed the Shapiro-Wilk test for normality, and groups were compared using two-tailed unpaired t-tests. Exact *p*-values are reported.

Taken together, these data demonstrate that *Crtap^-/-^* mice display disruptions in the development and postnatal maturation of tendons and ligaments. Moreover, the matrix disruptions caused by loss of CRTAP lead to the dysregulation of the postnatal tendon progenitor pool as well as tenocyte maturation.

### Collagen fibril formation is altered in heterozygous and Crtap^-/-^ mice

Tendons develop embryonically by increasing fibril length and number, whereas postnatal growth arises from an increase in fibril length and diameter – the latter of which is driven by the lateral fusion of smaller fibrils^4^. To investigate the role of Crtap in collagen fibril maturation, we utilized transmission electron microscopy (TEM) to examine changes in fibril diameter in flexor digitorum longus (FDL) and Achilles tendons as well as patellar ligaments (**Figure 3**). In the FDL tendon, there was a marked increase in small collagen fibrils (20-60 nm in size), a reduction in 80-320 nm fibrils, and a slight increase in larger fibrils (>340 nm in diameter) in *Crtap^-/-^* mice compared to wildtype (**Figure 3A,C,D**). Despite similarities seen in histology, heterozygous mutant FDL tendons also exhibited a slight increase in 20-40 nm fibrils in mice compared to wildtype controls (**Figure 3A-B,D**). Similar trends were observed for the Achilles tendon, namely an increase in small fibrils (20-60 nm), a reduction in 80-240 nm fibrils, and an increase in large fibrils (>280 nm) upon loss of *Crtap* (**Figure 3E,G-H**). In contrast to what we observed for the FDL, heterozygous Achilles tendons did not have increased numbers of smaller fibers (**Figure 3F,H**). Instead, a greater number of fibrils ranging from 140-200 nm in size were noted compared to wildtype controls.

**Figure 3.**
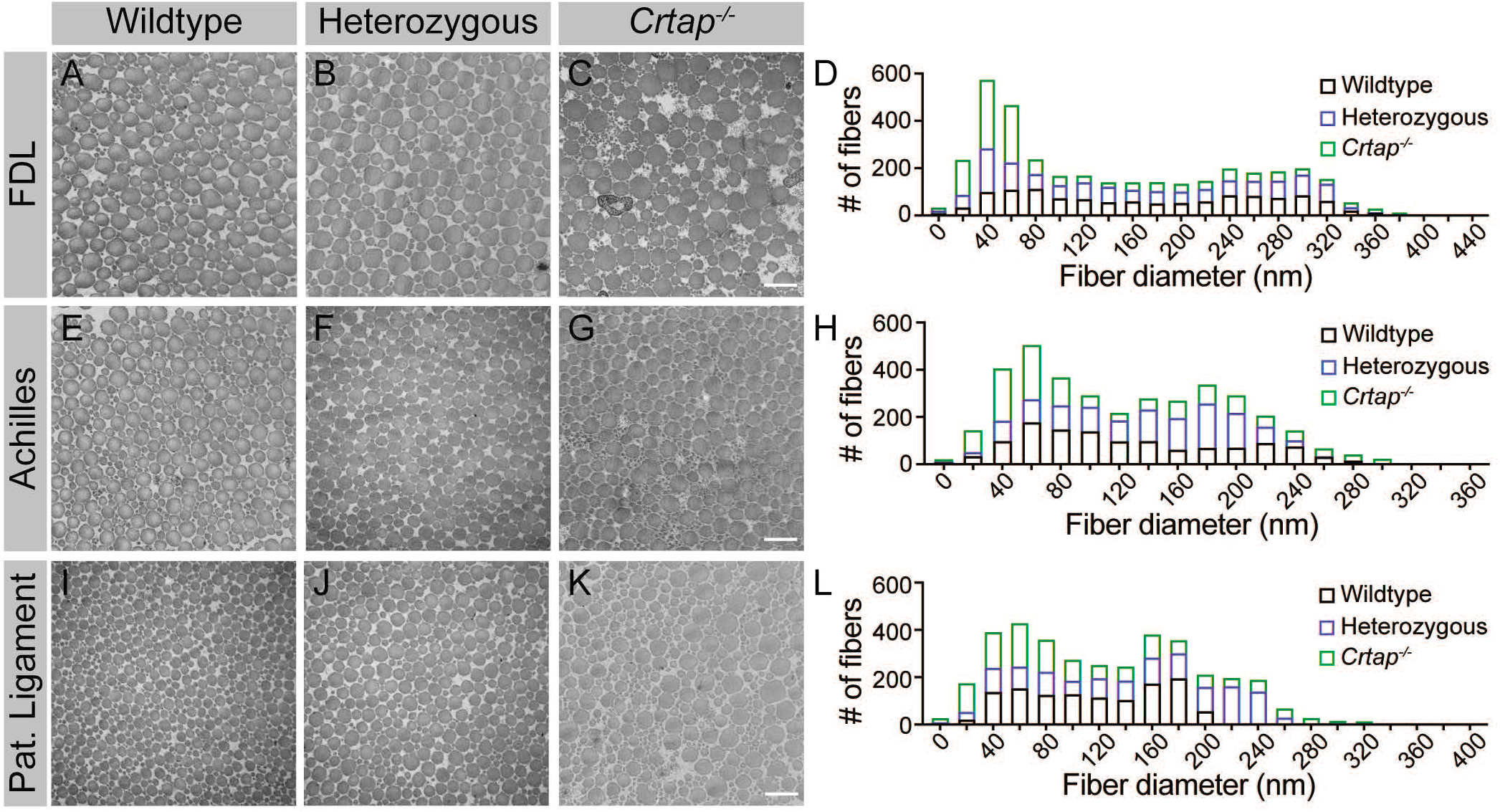
Collagen fibril diameter is altered in tendons and ligaments from heterozygous and *Crtap^-/-^* mice. **(A-C)** representative TEM images of 4-month FDL tendon collagen fibrils. Scale bar is 500-nm. **(D)** representative histogram of the size distribution for collagen fibrils in FDL tendons. Data are representative of n = 3 mice. **(E-G)** representative TEM images of 4-month Achilles tendon collagen fibrils. Scale bar is 500-nm. **(H)** representative histogram of the size distribution for collagen fibrils in Achilles tendons. Data are representative of n = 3 mice. **(I-K)** representative TEM images of 4-month patellar ligament collagen fibrils. Scale bar is 500-nm. **(L)** representative histogram of the size distribution for collagen fibrils in patellar ligaments. Data are representative of n = 3 mice per group.

Compared to the FDL and Achilles tendons, the greatest differences were seen within the patellar ligament, though the pattern of changes remained consistent (**Figure 3I-L**). Specifically, we observed a dramatic increase in 20 nm collagen fibrils compared to both heterozygous and wildtype mice (**Figure 3I-L**). Fibrils ranging from 100-180 nm in diameter were reduced in heterozygous and *Crtap^-/-^* mice compared to wildtype. Interestingly, the greatest difference from wildtype was an increase in large collagen fibrils (>200 nm) in both heterozygous and *Crtap^-/-^* mice (**Figure 3I-L**). Taken together, these data indicate that heterozygous and complete loss of CRTAP alter collagen fibril assembly in tendons and ligaments. In addition, the degree to which collagen assembly is affected is site-dependent.

### Collagen cross-linking is increased in heterozygous and Crtap^-/-^ mice

Along with P3H1 and CyPB, CRTAP is an integral part of the collagen prolyl 3-hydroxylation complex that is responsible for the 3-hydroxylation of Pro986 of the type I procollagen a1 chain^7^. Loss of this complex blocks 3-hydroxyproline formation and affects lysine hydroxylation and cross-linking in bone collagen^11, 12^; however, whether *Crtap^-/-^* tendons and ligaments display altered collagen cross-linking is unknown. To investigate this, we harvested tendons and ligaments at 1- and 4-months and assessed collagen cross-linking by quantifying the levels of hydroxylysyl-pyridinoline (HP) (**Figure 4**). Overall, we observed an increase in these stable, mature collagen cross-links from 1 to 4 months-of-age in all genotypes for the FDL and Achilles tendons (**Figure 4A-B**). In contrast, for the patellar ligament, age-dependent increases in collagen cross-links were only observed in *Crtap^-/-^* mice (**Figure 4C**). For FDL tendons, *Crtap^-/-^* mice had more of these collagen cross-links at 1- and 4-months compared to heterozygous and wildtype mice; however, the content of HP residues per collagen decreased with age in this tissue (**Figure 4A**). Interestingly, in Achilles tendons, an increase in collagen cross-linking was observed in both heterozygous and *Crtap^-/-^* mice at 1-month compared to wildtype. In contrast, at 4-months, only *Crtap^-/-^* mice had elevated collagen cross-links, and these levels were greater than those observed at the earlier time point (**Figure 4B**).

**Figure 4.**
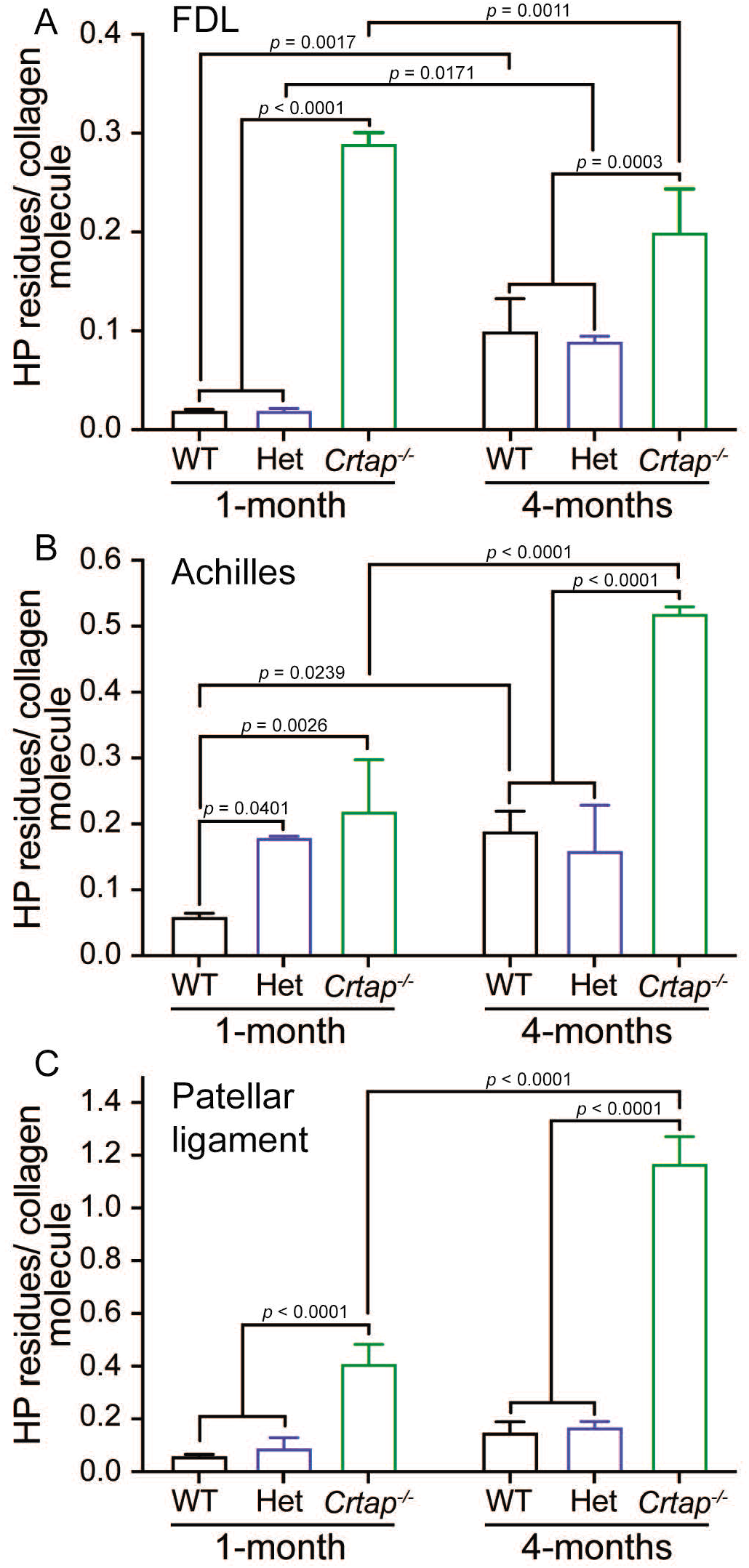
Collagen cross-linking is increased in tendons and ligaments from young and mature *Crtap^-/-^* mice. Quantification of collagen cross-links as hydroxylysyl-pyridinoline (HP) residues per collagen molecule for **(A)** FDL tendons; **(B)** Achilles tendons; and **(C)** patellar ligaments. Data are shown as means ± S.D. n = 3-4 mice per group. Data passed the Shapiro-Wilk test for normality, and groups were compared using one-way ANOVA with Tukey’s post-hoc tests. Exact *p*-values are reported.

Of the tissues examined, the patellar ligament showed the greatest increase in collagen cross-links both with time and across genotypes. Specifically, collagen cross-links were elevated by 5- to 10-fold in *Crtap^-/-^* patellar ligaments compared to heterozygous and wildtype at 1- and 4-months, respectively (**Figure 4C**). Taken together, these data suggest that *Crtap* is required for proper hydroxylation and cross-linking of collagen fibrils in tendons and ligaments in a semi-dominant fashion, as heterozygous mutant tendons/ligaments display a phenotype that is milder than the phenotype observed for homozygous mutant mice. Importantly, the chemical quality of collagen cross-linking appears to be spatiotemporally regulated and this regulation is differentially affected by the loss of a single or both copies of *Crtap*.

### Signaling and metabolic dysregulation in Crtap^-/-^ tendons and ligaments

To investigate the molecular changes underlying tendon/ligament phenotypes, we performed bulk RNA-seq with RNA isolated from Achilles tendons and patellar ligaments of 1-month-old wildtype and *Crtap^-/-^* mice. To determine global changes in differentially expressed genes as well as predicted upstream regulators, we performed Ingenuity Pathway Analysis (IPA, Qiagen, Germany). For the Achilles tendon, a total of 178 genes (consisting of 99 upregulated genes and 79 downregulated genes) were significantly differentially expressed between wildtype and *Crtap^-/-^* samples (**Figure 5A**). Of the top 30 differentially expressed genes, several extracellular matrix proteins, including matrilin-3 (*Matn3*), matrilin-4 (*Matn4*), and fibronectin 1 (*Fn1*), and proteolytic enzymes such as matrix metallopeptidase 2 (*Mmp2*) were dysregulated. Gene ontology analysis revealed “GO:000715 – Cell Adhesion”, “GO:0045778 – Positive Regulation of Ossification”, and “GO:0051928 – Positive Regulation of Calcium Ion Transport” to be enriched (**Figure 5B**). Examination of upstream regulators based on the differential gene expression data also revealed a predicted activation of TGF-β1 in *Crtap^-/-^* mice and predicted inhibition of dystrophin (DMD) along with several for which activation state was unclear, including platelet-derived growth factor-BB (PDGF-BB), β-catenin (CTNNB1), and tumor necrosis factor (TNF) (**Figure 5C**).

**Figure 5.**
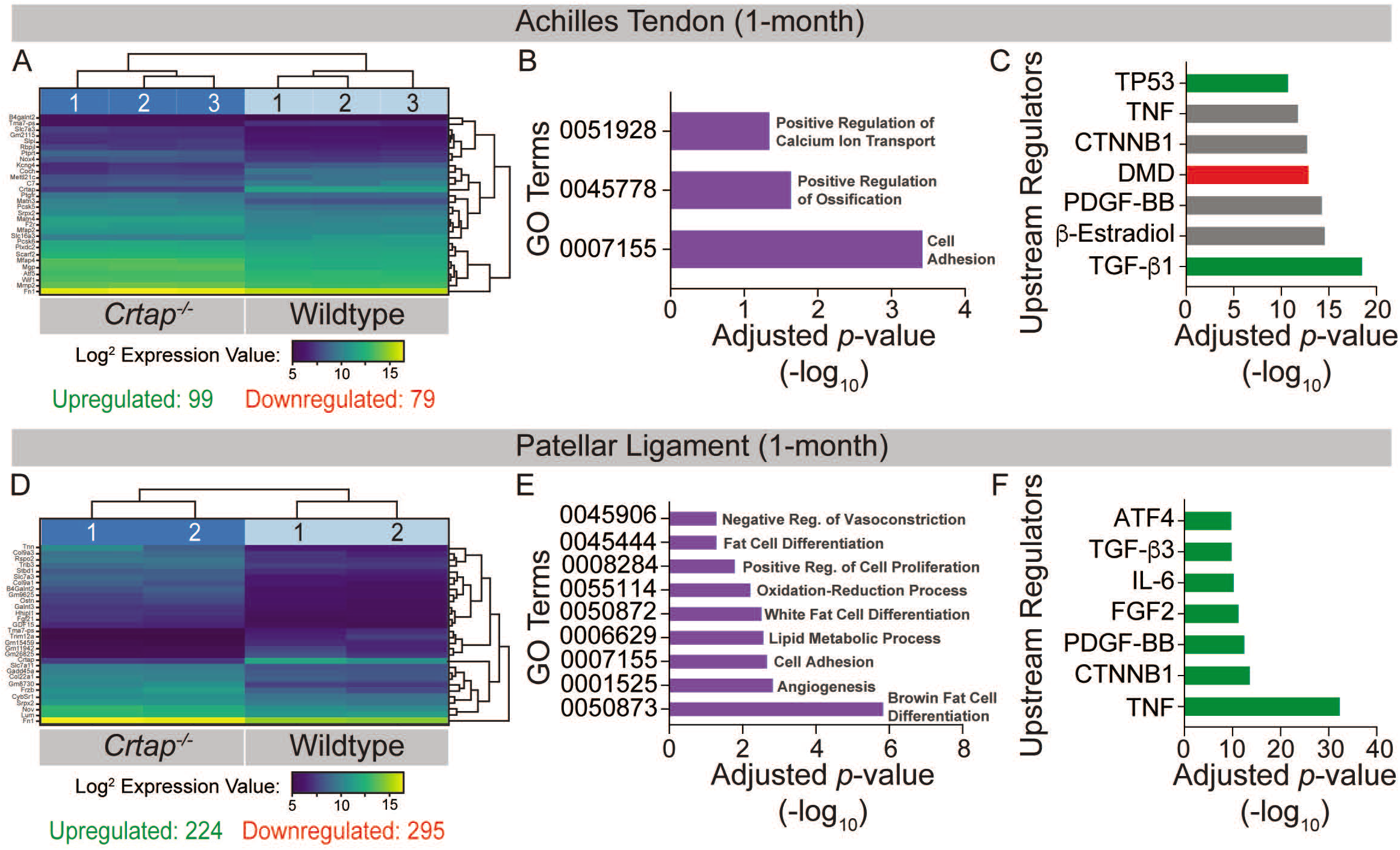
Transcriptome analysis of tendons and ligaments from 1-month-old *Crtap^-/-^* mice. DESeq2 was used to compare gene expression between wildtype and *Crtap^-/-^* Achilles tendon and patellar ligament RNA samples, and genes with an adjusted *p*-value < 0.05 and absolute log_2_ fold change > 1 were considered as differentially expressed. **(A)** a bi-clustering heatmap of the top 30 differentially expressed genes between wildtype and *Crtap^-/-^* Achilles tendons sorted by adjusted *p*-value and plotted according to log_2_ transformed expression values. The Wald test was used to generate p-values and log_2_ fold changes. **(B)** significantly differentially expressed genes between Achilles tendons from wildtype and *Crtap^-/-^* mice were clustered by their gene ontology, and enrichment for gene ontology terms was tested using Fisher exact test. All gene ontology terms with an adjusted *p*-value < 0.05 are plotted according to their −log_10_ adjusted *p*-value. **(C)** select upstream regulators predicted as being activated (shown in green), inhibited (shown in red), or of unclear state (shown in grey) in *Crtap^-/-^* compared to wildtype Achilles tendons plotted according to their −log_10_ adjusted *p*-value. n = 3 mice per genotype for **(A-C)**. **(D)** a bi-clustering heatmap of the top 30 differentially expressed genes between wildtype and *Crtap^-/-^* patellar ligaments sorted by adjusted *p*-value and plotted according to log_2_ transformed expression values. The Wald test was used to generate p-values and log_2_ fold changes. **(E)** significantly differentially expressed genes between Achilles tendons from wildtype and *Crtap^-/-^* mice were clustered by their gene ontology, and enrichment for gene ontology terms was tested using Fisher exact test. All gene ontology terms with an adjusted *p*-value < 0.05 are plotted according to their −log_10_ adjusted *p*-value. **(F)** select upstream regulators predicted as being activated (shown in green), inhibited (shown in red), or of unclear state (shown in grey) in *Crtap^-/-^* compared to wildtype patellar ligaments plotted according to −log_10_ adjusted *p*-value. n = 2 mice per genotype for **(D-F)**.

In keeping with the increased severity seen in ligaments from *Crtap^-/-^* mice compared to tendons, a greater number of total genes were differentially expressed between wildtype and *Crtap^-/-^* patellar ligament samples (**Figure 5D**). Specifically, we saw significant differential expression of 519 genes, with 224 being upregulated and 295 downregulated in *Crtap^-/-^* compared to wildtype. Several of the top 30 differentially expressed genes were small collagens such as type IX collagen α3 chain (*Col9a3*), type IX collagen α1 chain (*Col9a1*), and type XXII collagen α1 chain (*Col22a1*) as well as other extracellular matrix proteins, including lumican (Lum) and fibronectin 1 (*Fn1*). Unlike the Achilles tendon, gene ontology analysis revealed significant enrichment for metabolic processes, including “GO:0055114 – Oxidation-Reduction Process”, “GO:0050873 – Brown Fat Cell Differentiation”, and “GO:0050872 – White Fat Cell Differentiation” as well as for “GO:0008284 – Positive Regulation of Proliferation” and “GO:0001525 – Angiogenesis” (**Figure 5E**). Examination of upstream regulators also suggested a highly significant activation of TNF and interleukin 6 (IL-6) as well as fibroblast growth factor 2 (FGF2) and TGF-β3 in *Crtap^-/-^* patellar ligaments (**Figure 5F**). Indeed, the representation of TNF, TGF-β, and PDGF-BB in the ligament dataset is consistent with but more severe than that observed for those same regulators in the Achilles tendon results (**Figure 5C**).

### Loss of CRTAP leads to deficiencies in motor activity, coordination, and strength

To investigate whether the tendon phenotypes observed in *Crtap^-/-^* mice have functional consequences, we performed a series of behavioral assays at 4 months-of-age. Using the open field assay to quantify changes in spontaneous motor activity, we observed that *Crtap^-/-^* mice displayed significant reductions in both horizontal and vertical activity compared to heterozygous and wildtype mice (**Figure 6A-B**). We next examined changes in motor coordination and endurance using the rotarod assay. While no genotype-dependent differences were observed during the learning phase of the assessment (Trials 1-5), *Crtap^-/-^* displayed a reduction in latency to fall for Trials 6, 9-10 compared to wildtype and Trials 6, 8-10 compared to heterozygous mice (**Figure 6C**). To confirm this observation, we evaluated the mice using the grid foot slip assay – an alternative metric for motor coordination. In this regard, we found that *Crtap^-/-^* mice exhibited a modest increase in forelimb and hindlimb foot slips compared to heterozygous and wildtype mice (**Figure 6D-E**). Taken together, these findings indicate that *Crtap^-/-^* mice have deficiencies in motor activity and coordination compared to controls.

**Figure 6.**
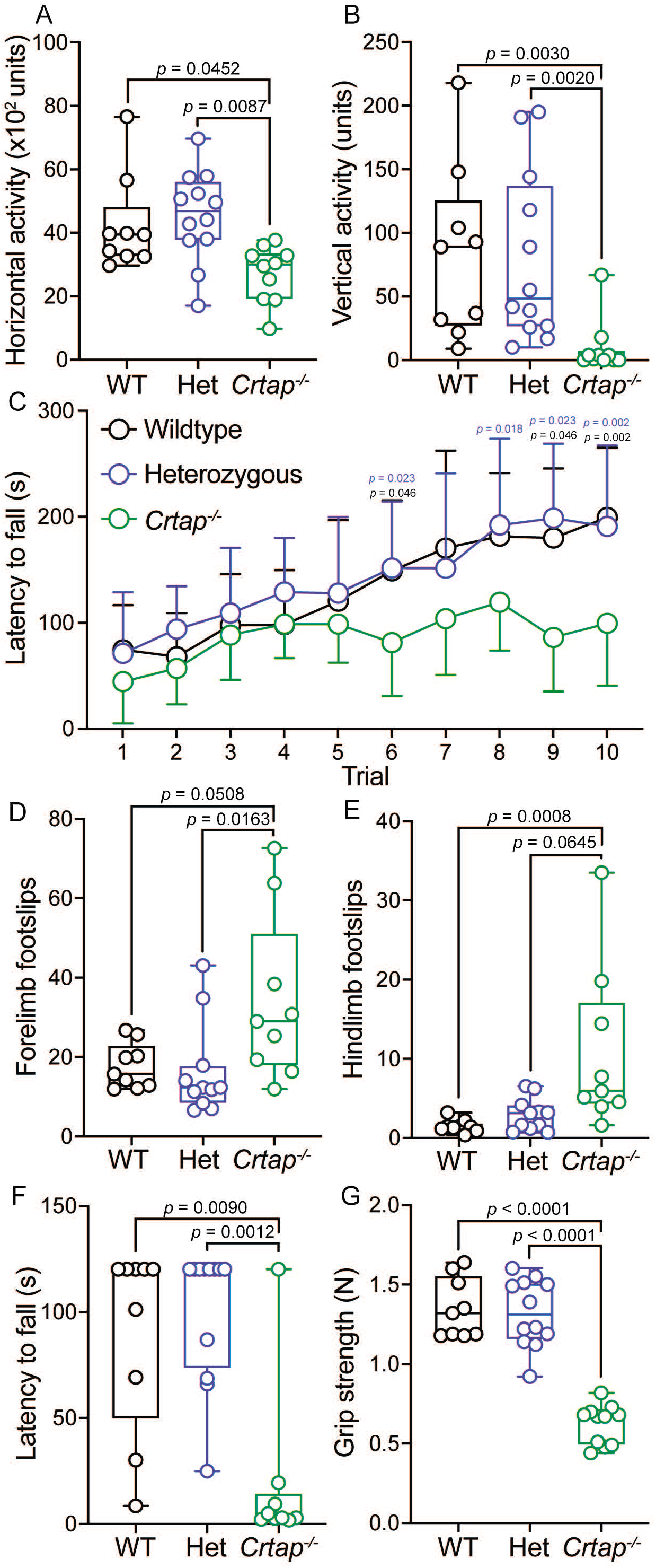
Motor activity and coordination are impaired in 4-months-old *Crtap^-/-^* mice. **(A-B)** quantification of spontaneous motor activity including horizontal **(A)** and vertical **(B)** activity over a 30-min period using the open field assay. Data are min-to-max box and whisker plots with individual points indicated. n = 9-12 mice per group. For **(A)**, data passed the Shapiro-Wilk test for normality, and groups were compared using one-way ANOVA with Tukey’s post-hoc tests. For **(B)**, data failed the Shapiro-Wilk test for normality, and groups were compared using Kruskal-Wallis with Dunn’s post-hoc tests. Exact p-values are reported for **(A-B)**. **(C)** quantification of motor activity, coordination, and endurance across 10 trials conducted over 2 days using an accelerating rotarod assay. Data are means ± S.D. n = 9-12 mice per group. Groups were compared using repeated-measures two-way ANOVA with Tukey’s post-hoc tests. Exact *p*-values are reported, with black *p*-values being compared to wildtype, and blue *p*-values being compared to heterozygous mice. **(D-E)** quantification of forelimb **(D)** and hindlimb **(E)** motor coordination using the grid foot slip assay. Data are min-to-max box and whisker plots with individual points indicated. n = 9-11 mice per group. Data failed the Shapiro-Wilk test for normality, and groups were compared using Kruskal-Wallis with Dunn’s post-hoc tests. Exact p-values are reported. **(F)** quantification of forelimb and hindlimb grip strength using the inverted grid assay conducted for 120 s. A reduction in latency to fall indicates reduced strength. Data are min-to-max box and whisker plots with individual points indicated. n = 8-9 mice per group. Data failed the Shapiro-Wilk test for normality, and groups were compared using Kruskal-Wallis with Dunn’s post-hoc tests. Exact p-values are reported. **(G)** quantification of forelimb grip strength in N of force measured over 3 trials and then averaged. Data are min-to-max box and whisker plots with individual points indicated. n = 9-12 mice per group. Data passed the Shapiro-Wilk test for normality, and groups were compared using one-way ANOVA with Tukey’s post-hoc tests. Exact *p*-values are reported.

We next evaluated strength in the *Crtap^-/-^* mice using the inverted grid and grip strength assays. Interestingly, we observed a decrease in the latency to fall during the inverted grid assay for *Crtap^-/-^* mice compared to heterozygous and wildtype controls (**Figure 6F**). Using a more quantitative metric, we examined these mice using the grip strength test and found that while wildtype and heterozygous mice could generate approximately 1.3 N of force, mice lacking CRTAP were weaker with a mean grip strength of 0.62 N (**Figure 6G**). Thus, *Crtap^-/-^* mice display significant reductions in strength together with perturbations in motor activity and coordination – behavioral changes that could be related in part to their tendon phenotype.

## DISCUSSION

In this study, we examined the histological, ultrastructural, biochemical, and transcriptional characteristics of tendons and ligaments in the *Crtap^-/-^* mouse model of severe, recessive OI. We demonstrate that at 1- and 4 months-of-age, *Crtap^-/-^* have thinner, more cellular Achilles tendons and patellar ligaments with a reduction in tendon progenitors compared to heterozygous and wildtype mice. Using phase-contrast μCT imaging, we confirmed that tissue volume is reduced in *Crtap^-/-^* patellar ligaments at 4-months than those from heterozygous and wildtype mice. Examining collagen fibril organization, we found a marked alteration in small and large fibrils in both heterozygous and *Crtap^-/-^* mice compared to wildtype that varied in severity depending on the tendon or ligament being examined. Stable, HP cross-links were also elevated at 1- and 4-months in *Crtap^-/-^* mice compared to wildtype, which indicates an increase in telopeptide lysine hydroxylation and irreversible intermolecular cross-links (see below). RNA-seq analysis revealed alterations in extracellular matrix proteins, growth factor signaling, and metabolic pathways in *Crtap^-/-^* tendons and ligaments compared to wildtype controls. Finally, we demonstrate that *Crtap^-/-^* mice exhibit motor impairments concomitant with reductions in grip strength – a phenomenon that may be related to the tendon pathology observed in these mice.

Collagen ultrastructure and cross-linking has been well-documented in the bones of mouse models for both dominant and recessive OI; however, quantitative differences in the volume of tendons and ligaments in these models has not been examined and tendon/ligament histological characterization has been modest. In this regard, we demonstrate that *Crtap^-/-^* mice have thinner Achilles tendons and patellar ligaments at 1- and 4-months. Despite the reduction in size and extracellular matrix, tendons and ligaments from these mice are hypercellular compared to wildtype and heterozygous animals. At the same time, there is a significant reduction in postnatal CD146+CD200+ tendon progenitors with a concomitant increase of immature cells. While the alterations in tendon and ligament size and cellularity may be a result of alterations in the collagen extracellular matrix, it could also be associated with alterations in cellular signaling. In this regard, TGF-β signaling is upregulated in bones from *Crtap^-/-^* mice^34^, and elevated TGF-β signaling has been noted in mouse models with increased tendon cellularity and alterations in collagen fibril distribution^35^. Indeed, our RNA-seq analysis identified TGF-β signaling as an activated upstream regulator predicted to be driving differential gene expression observed in the tendons and ligaments from *Crtap^-/-^* mice. We also observed significant metabolic dysregulation in *Crtap^-/-^* ligaments. Interestingly, this a phenotype that has been observed in mammalian target of rapamycin complex 1 (mTORC1) knockout mouse tendons that have a similar histological phenotype to those of *Crtap^-/-^* mice^35^. These data are consistent with the principle that an altered extracellular matrix can lead to altered cellular signaling as previously shown for vascular tissue in Marfan syndrome and OI bone. Here, we extend this theme to tendon and ligamentous defects in OI.

Collagen fibril assembly is a dynamic process that begins with the formation of many small fibrils that grow longitudinally during development, followed by appositional fusion and continued longitudinal growth as tendons mature^4^. In the present study, we found an increased proportion of small and large collagen fibrils in the FDL and Achilles tendons as well as patellar ligaments of *Crtap^-/-^* mice compared to wildtype. These distributions varied in severity across the three tissues, with the greatest differences being observed in the patellar ligament. *Lepre1^-/-^* mice have been reported to display an increased proportion of small collagen fibrils in tail tendons^19^, indicating that loss of the 3-prolyl hydroxylase complex alters collagen fibrillogenesis. At the same time, the increased number of small fibers was more pronounced in *Lepre1^-/-^* mice, and there was no evidence of an increased proportion of large fibers^19^ as observed for *Crtap^-/-^* mice in this study. Similar to the *Lepre1^-/-^* mouse model, mice lacking CypB also exhibit a pronounced increase in the number of small collagen fibrils within tail tendons^18^. Together, this suggests that despite forming a complex with P3h1 and CypB, loss of Crtap has distinct consequences on collagen fibrillogenesis. In this regard, it is important to note that tail tendons were used for collagen fibril distribution assessments in both *Lepre1^-/-^* and *Ppib^-/-^* mice^18, 19^, whereas appendicular tendons and ligaments were examined in our study. The difference in tissue type examined might also explain why we observed unique alterations in collagen fibril distribution in heterozygous *Crtap* mice, whereas *CypB* heterozygous tendons were indistinguishable from wildtype^18^.

Type I procollagen molecules undergo post-translational modifications within the endoplasmic reticulum, including lysyl-hydroxylation and prolyl-hydroxylation, that are critical for proper collagen synthesis, transport, and stability. Specifically, telopeptide lysine hydroxylation results in mature lysyl-pyridinoline (LP) or hydroxylysyl-pyridinoline (HP) residues after lysyl oxidase oxidation, which as permanent, irreversible crosslinks play a role in regulating fibril growth and strength^25, 36^. Previous literature has demonstrated that loss of the 3-prolyl hydroxylase complex caused by deletion of P3h1 (*Lepre1^-/-^*) or Crtap (*Crtap^-/-^*) prevents prolyl 3-hydroxylation of clade A (type I, II and III) collagens and can lead to changes in lysine post-translational modifications due to loss of its chaperone function^10, 11^. In this study, we found that mature collagen cross-links (HP residues per collagen) are markedly increased in the FDL and Achilles tendons as well as the patellar ligament of *Crtap^-/-^* mice relative to wildtype. Interestingly, we observed increased cross-links in the Achilles, but not FDL or patellar tendons of 1-month old heterozygous mice, indicating a mild haploinsufficient effect of Crtap on this tendon biochemical property thereby suggesting a rate-limited contribution of this complex in this function. Outside of the genotype-specific effects, we saw an age-dependent increase in HP residues per collagen in all genotypes. This observation is consistent with a study by Taga and colleagues that reported an increase in 3-hydroxyproline residues in rat tendon collagen (but not bone or skin) that plateaued at 3 months-of-age^37^. Together with the TEM analyses, these data suggest that altered collagen cross-linking in tendons and ligaments from *Crtap^-/-^* mice may adversely affect collagen fibril assembly.

In addition to skeletal deformities and frequent fractures, severe OI is associated with motor impairments including gait abnormalities, chronic pain, and reduced muscle strength^8, 9, 38^. In this study, we showed that *Crtap^-/-^* mice exhibit reduced motor activity and coordination using the open field, rotarod, and grid foot slip assays. We also observed a reduction in latency to fall on the inverted grid assay that was mirrored by a dramatic loss of grip strength compared to heterozygous and wildtype mice. These results are consistent with findings reported for the *Col1a1^Jrt/+^* mouse model of severe OI and Ehlers-Danlos syndrome^17^. Specifically, Abdelaziz and colleagues found that *Col1a1^Jrt/+^* mice displayed reduced motor activity using the open field and running wheel assays – a phenotype they attributed to thermal hyperalgesia and mechanical allodynia in these mice^39^. In this regard, the reduced vertical activity we observed in *Crtap^-/-^* mice may be indicative of a pain or spinal phenotype, suggesting that the characterization of pain in this model is a useful avenue for future research. Overall, this study represents one of the first extensive characterizations of behavioral deficits in a mouse model of severe, recessive OI that also correlates these deficits to tendon phenotypes.

Taken together, this study provides the first evidence for tendon and ligament phenotypes in the *Crtap^-/-^* mouse model of severe recessive OI. We also provide compelling evidence for a strong motor activity and coordination phenotype in these mice. As the quality of life is so impacted in patients with OI, a more comprehensive evaluation of behavioral outcomes in future preclinical studies may provide important insights into the efficacy of therapeutic interventions.

## AUTHOR CONTRIBUTIONS

M. W. Grol: Conception and design of the study, acquisition, analysis and interpretation of data, drafting and editing of the manuscript

N. A. Haelterman: Acquisition and analysis of data, editing of the manuscript

J. Lim: Acquisition and analysis of data, editing of the manuscript

E. Munivez: Acquisition of data

M. Archer: Acquisition and analysis of data

D. M. Hudson: Acquisition, analysis, and interpretation of data, editing of the manuscript

S. F. Tufa: Acquisition and analysis of data

D. R. Keene: Acquisition, analysis, and interpretation of data, editing of the manuscript

K. Lei: Acquisition and analysis of data

D. Park: Analysis and interpretation of data

D. R. Eyre: Analysis and interpretation of data, editing of the manuscript

B H. Lee: Conception and design of the study, interpretation of data, editing of the manuscript

## REFERENCES

1. Nourissat G, Berenbaum F, Duprez D. Tendon injury: from biology to tendon repair. Nat Rev Rheumatol 2015; 11: 223–233.

2. Screen HR, Berk DE, Kadler KE, Ramirez F, Young MF. Tendon functional extracellular matrix. J Orthop Res 2015; 33: 793–799.

3. Kannus P. Structure of the tendon connective tissue. Scand J Med Sci Sports 2000; 10: 312–320.

4. Kalson NS, Lu Y, Taylor SH, Starborg T, Holmes DF, Kadler KE. A structure-based extracellular matrix expansion mechanism of fibrous tissue growth. Elife 2015; 4.

5. Saito M, Marumo K. Collagen cross-links as a determinant of bone quality: a possible explanation for bone fragility in aging, osteoporosis, and diabetes mellitus. Osteoporos Int 2010; 21: 195–214.

6. Alford AI, Kozloff KM, Hankenson KD. Extracellular matrix networks in bone remodeling. Int J Biochem Cell Biol 2015; 65: 20–31.

7. Lim J, Grafe I, Alexander S, Lee B. Genetic causes and mechanisms of osteogenesis imperfecta. Bone 2017; 102: 40–49.

8. Arponen H, Makitie O, Waltimo-Siren J. Association between joint hypermobility, scoliosis, and cranial base anomalies in paediatric osteogenesis imperfecta patients: a retrospective cross-sectional study. BMC Musculoskelet Disord 2014; 15: 428.

9. Primorac D, Anticevic D, Barisic I, Hudetz D, Ivkovic A. Osteogenesis imperfecta--multi-systemic and life-long disease that affects whole family. Coll Antropol 2014; 38: 767–772.

10. Hudson DM, Eyre DR. Collagen prolyl 3-hydroxylation: a major role for a minor post-translational modification? Connect Tissue Res 2013; 54: 245–251.

11. Morello R, Bertin TK, Chen Y, Hicks J, Tonachini L, Monticone M, et al. CRTAP is required for prolyl 3-hydroxylation and mutations cause recessive osteogenesis imperfecta. Cell 2006; 127: 291–304.

12. Baldridge D, Schwarze U, Morello R, Lennington J, Bertin TK, Pace JM, et al. CRTAP and LEPRE1 mutations in recessive osteogenesis imperfecta. Hum Mutat 2008; 29: 1435–1442.

13. Barnes AM, Chang W, Morello R, Cabral WA, Weis M, Eyre DR, et al. Deficiency of cartilage-associated protein in recessive lethal osteogenesis imperfecta. N Engl J Med 2006; 355: 2757–2764.

14. Cabral WA, Chang W, Barnes AM, Weis M, Scott MA, Leikin S, et al. Prolyl 3-hydroxylase 1 deficiency causes a recessive metabolic bone disorder resembling lethal/severe osteogenesis imperfecta. Nat Genet 2007; 39: 359–365.

15. van Dijk FS, Nesbitt IM, Zwikstra EH, Nikkels PG, Piersma SR, Fratantoni SA, et al. PPIB mutations cause severe osteogenesis imperfecta. Am J Hum Genet 2009; 85: 521–527.

16. Baldridge D, Lennington J, Weis M, Homan EP, Jiang MM, Munivez E, et al. Generalized connective tissue disease in *Crtap^-/-^* mouse. PLoS One 2010; 5: e10560.

17. Chen F, Guo R, Itoh S, Moreno L, Rosenthal E, Zappitelli T, et al. First mouse model for combined osteogenesis imperfecta and Ehlers-Danlos syndrome. J Bone Miner Res 2014; 29: 1412–1423.

18. Terajima M, Taga Y, Chen Y, Cabral WA, Hou-Fu G, Srisawasdi S, et al. Cyclophilin-B modulates collagen cross-linking by differentially affecting lysine hydroxylation in the helical and telopeptidyl domains of tendon type I collagen. J Biol Chem 2016; 291: 9501–9512.

19. Vranka JA, Pokidysheva E, Hayashi L, Zientek K, Mizuno K, Ishikawa Y, et al. Prolyl 3-hydroxylase 1 null mice display abnormalities in fibrillar collagen-rich tissues such as tendons, skin, and bones. J Biol Chem 2010; 285: 17253–17262.

20. Schindelin J, Arganda-Carreras I, Frise E, Kaynig V, Longair M, Pietzsch T, et al. Fiji: an open-source platform for biological-image analysis. Nat Methods 2012; 9: 676–682.

21. Nixon AJ, Grol MW, Lang HM, Ruan MZC, Stone A, Begum L, et al. Disease-modifying osteoarthritis treatment with interleukin-1 receptor antagonist gene therapy in small and large animal models. Arthritis Rheumatol 2018; 70: 1757–1768.

22. Ruan MZ, Erez A, Guse K, Dawson B, Bertin T, Chen Y, et al. Proteoglycan 4 expression protects against the development of osteoarthritis. Sci Transl Med 2013; 5: 176ra134.

23. Ruan MZ, Patel RM, Dawson BC, Jiang MM, Lee BH. Pain, motor and gait assessment of murine osteoarthritis in a cruciate ligament transection model. Osteoarthritis Cartilage 2013; 21:1355–1364.

24. Stone A, Grol MW, Ruan MZC, Dawson B, Chen Y, Jiang MM, et al. Combinatorial *Prg4* and *Il-1ra* gene therapy protects against hyperalgesia and cartilage degeneration in post-traumatic osteoarthritis. Hum Gene Ther 2019; 30: 225–235.

25. Eyre D. Collagen cross-linking amino acids. Methods Enzymol 1987; 144: 115–139.

26. Hudson DM, Archer M, King KB, Eyre DR. Glycation of type I collagen selectively targets the same helical domain lysine sites as lysyl oxidase-mediated cross-linking. J Biol Chem 2018; 293: 15620–15627.

27. Steinmann S, Pfeifer CG, Brochhausen C, Docheva D. Spectrum of tendon pathologies: triggers, trails and end-state. Int J Mol Sci 2020; 21.

28. Dyment NA, Hagiwara Y, Matthews BG, Li Y, Kalajzic I, Rowe DW. Lineage tracing of resident tendon progenitor cells during growth and natural healing. PLoS One 2014; 9: e96113.

29. Dyment NA, Liu CF, Kazemi N, Aschbacher-Smith LE, Kenter K, Breidenbach AP, et al. The paratenon contributes to scleraxis-expressing cells during patellar tendon healing. PLoS One 2013; 8: e59944.

30. Howell K, Chien C, Bell R, Laudier D, Tufa SF, Keene DR, et al. Novel model of tendon regeneration reveals distinct cell mechanisms underlying regenerative and fibrotic tendon healing. Sci Rep 2017; 7: 45238.

31. Harvey T, Flamenco S, Fan CM. A Tppp3(+)Pdgfra(+) tendon stem cell population contributes to regeneration and reveals a shared role for PDGF signalling in regeneration and fibrosis. Nat Cell Biol 2019; 21: 1490–1503.

32. Lee CH, Lee FY, Tarafder S, Kao K, Jun Y, Yang G, et al. Harnessing endogenous stem/progenitor cells for tendon regeneration. J Clin Invest 2015; 125: 2690–2701.

33. Feng H, Xing W, Han Y, Sun J, Kong M, Gao B, et al. Tendon-derived cathepsin K-expressing progenitor cells activate Hedgehog signaling to drive heterotopic ossification. J Clin Invest 2020.

34. Grafe I, Yang T, Alexander S, Homan EP, Lietman C, Jiang MM, et al. Excessive transforming growth factor-beta signaling is a common mechanism in osteogenesis imperfecta. Nat Med 2014; 20: 670–675.

35. Lim J, Munivez E, Jiang MM, Song IW, Gannon F, Keene DR, et al. mTORC1 Signaling is a critical regulator of postnatal tendon development. Sci Rep 2017; 7: 17175.

36. Bateman JF, Boot-Handford RP, Lamande SR. Genetic diseases of connective tissues: cellular and extracellular effects of ECM mutations. Nat Rev Genet 2009; 10: 173–183.

37. Taga Y, Kusubata M, Ogawa-Goto K, Hattori S. Developmental stage-dependent regulation of prolyl 3-hydroxylation in tendon type I collagen. J Biol Chem 2016; 291: 837–847.

38. Garman CR, Graf A, Krzak J, Caudill A, Smith P, Harris G. Gait deviations in children with osteogenesis imperfecta type I. J Pediatr Orthop 2019; 39: e641–e646.

39. Abdelaziz DM, Abdullah S, Magnussen C, Ribeiro-da-Silva A, Komarova SV, Rauch F, et al. Behavioral signs of pain and functional impairment in a mouse model of osteogenesis imperfecta. Bone 2015; 81: 400–406.

